## Supplementary Material for "No evidence for increased transmissibility from recurrent mutations in SARS-CoV-2"

Damien Richard^2,3*^

Cedric CS. Tan^1^

Liam P. Shaw^4^

Mislav Acman^1^

François Balloux^1+^

^1^ UCL Genetics Institute, University College London, London WC1E 6BT, UK

^2^ Cirad, UMR PVBMT, F-97410 St Pierre, Réunion, France

^3^ Université de la Réunion, UMR PVBMT, F-97490 St Denis, Réunion, France

^4^ Nuffield Department of Medicine, John Radcliffe Hospital, University of Oxford, Oxford OX3 9DU, UK

*contributed equally

+ corresponding; (Lucy van Dorp) and (François Balloux)

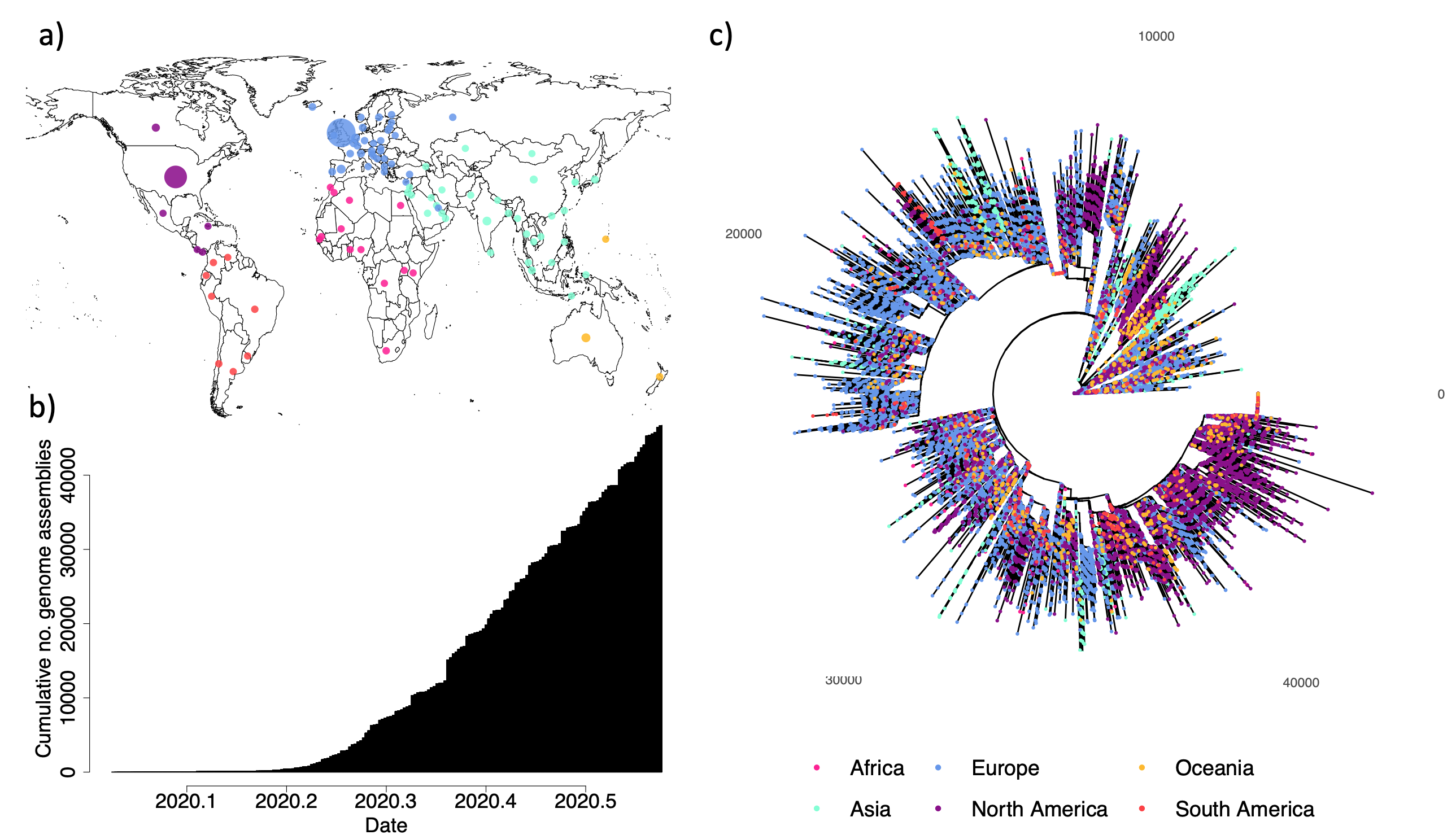
**Figure S1:** a) Geographic origin of sequences included in phylogenetic analysis (downloaded 30/07/2020). Purple: North America, coral: South America, blue: Europe, pink: Africa, green: Asia, Oceania: orange. Circles are proportional to the number of uploaded assemblies. b) Date of sample collection of all assemblies included in the dataset. c) Maximum likelihood phylogenetic tree (radial format). Tips are annotated by location of sampling as given in panel a).

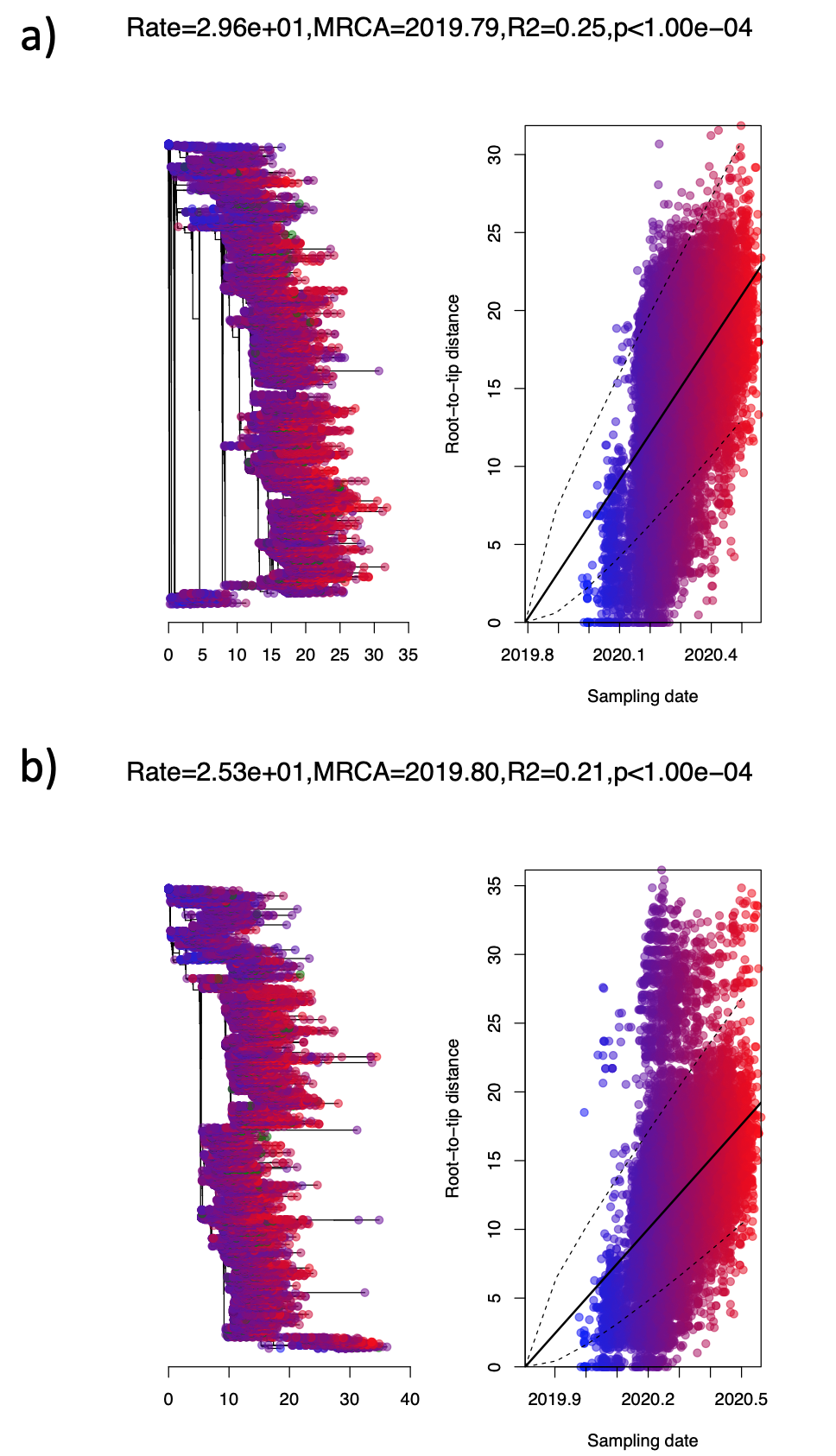

**Figure S2:** Root-to-tip regression performed using the *roottotip()* function provided by BactDating (see main text Methods) for both masked alignments (a- de Maio *et al.*, b – NextStrain). X-axis provides the time of sampling with y providing the root-to-tip distance across the rooted maximum likelihood phylogenetic tree. BactDating provides the *p*-value following 10,000 permutations of the tree sampling dates.

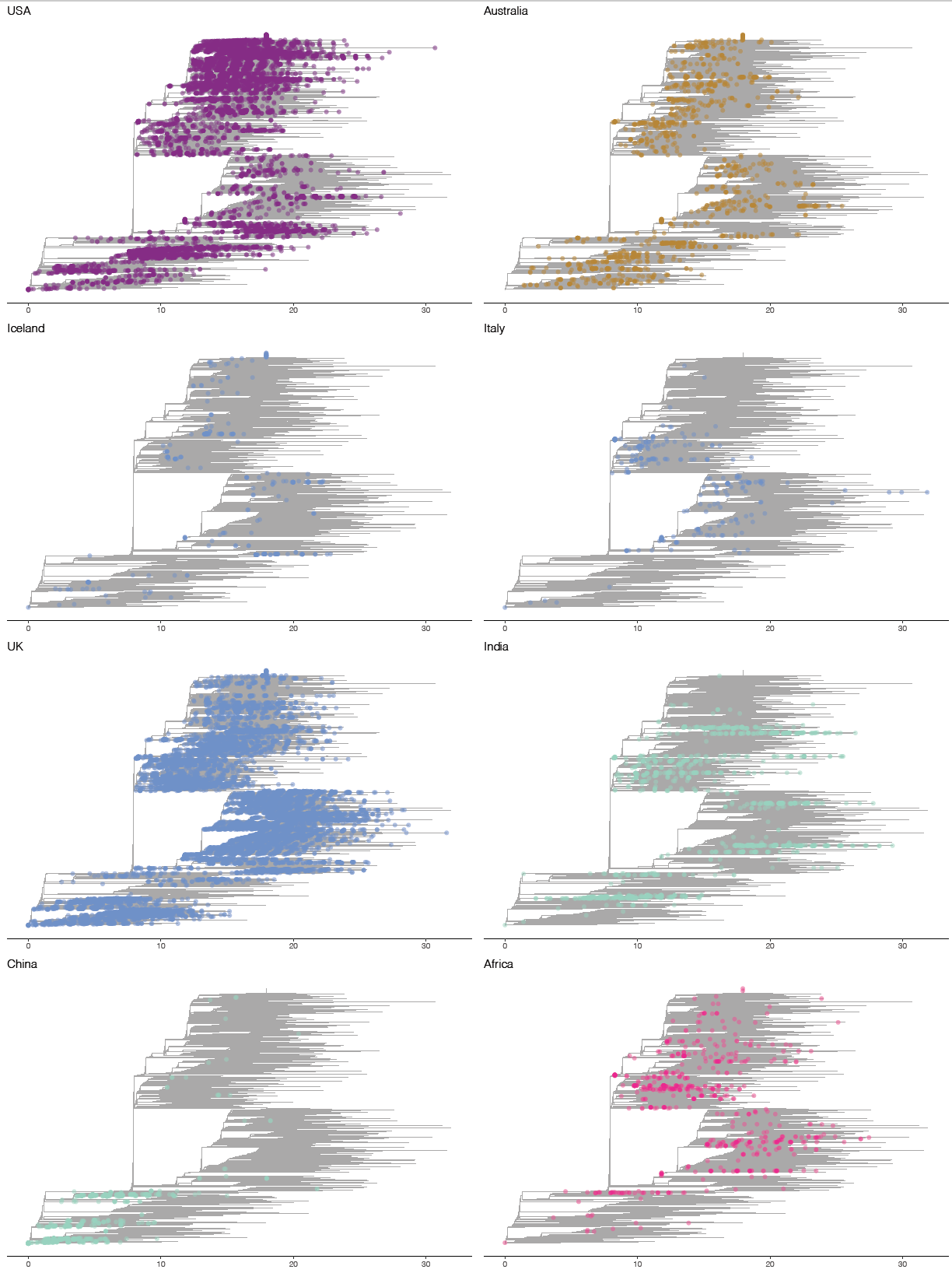

**Figure S3**: Maximum likelihood phylogenetic trees rooted on Wuhan-Hu-1 built using the Augur tree rapid phylodynamic pipeline. Tips are coloured by regions as given in main text Figure 1 and Figure S1. Genomes sampled in specific geographic regions are highlighted as indicated on the top-left of each plot.

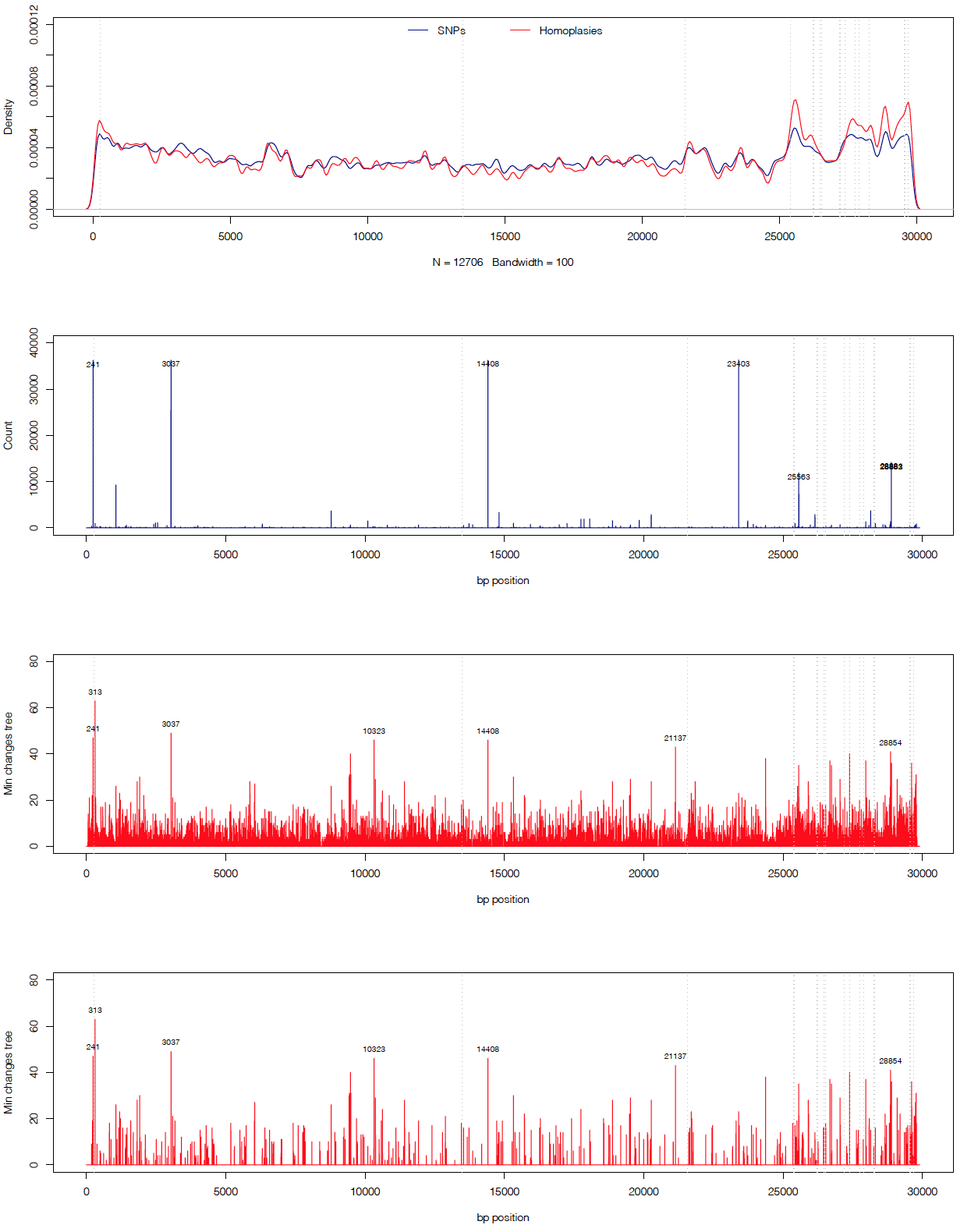

**Figure S4:** Genome-wide distribution of SNPs and homoplasies in SARS-CoV-2. Top provides the density of raw inferred SNPs and homoplasies genome-wide. The SNP count is provided with SNPs occurring in >10,000 isolates annotated. The raw count of 5,710 homoplasies is given in red with those responsible for >40 minimum changes on the tree annotated. This is filtered to a final set of 398 recurrent mutations (bottom panel), again annotated for those contributing to >40 minimum changes on the tree. A full list of filtered and non-filtered homoplasies for this alignment is provided in **Supplementary Table S3.**

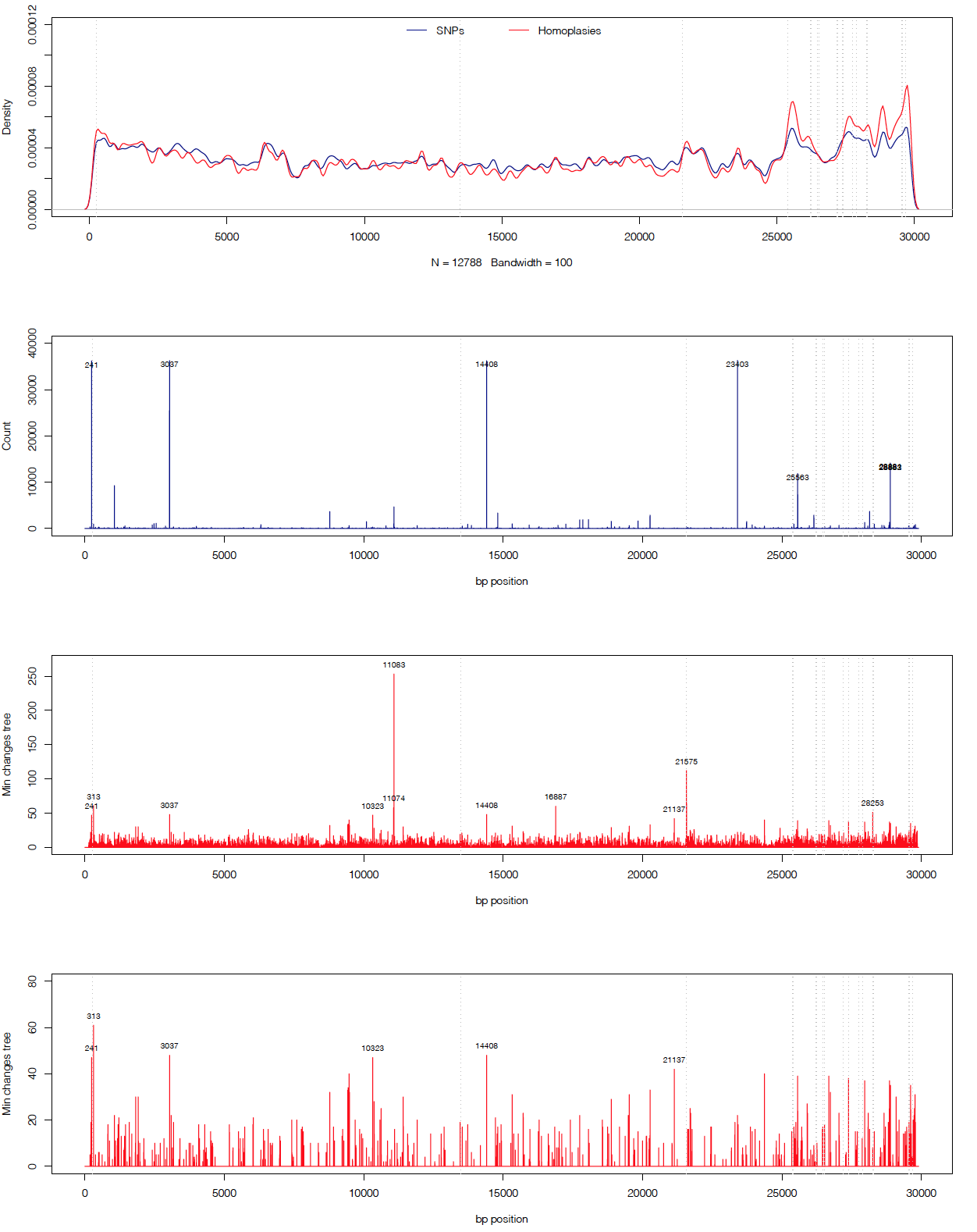
**Figure S5:** Genome-wide distribution of SNPs and homoplasies in SARS-CoV-2 following the NextStrain masking strategy (see **Supplementary Table S5**). Top provides the density of raw inferred SNPs and homoplasies genome-wide. The SNP count is provided with SNPs occurring in >10,000 isolates annotated. The raw count of 5,793 homoplasies is given in red with those responsible for >40 minimum changes on the tree annotated. This is filtered to a final set of 411 recurrent mutations (bottom panel), again annotated for those contributing to >40 minimum changes on the tree. A full list of filtered and non-filtered homoplasies for this alignment is provided in **Supplementary Table S3.**

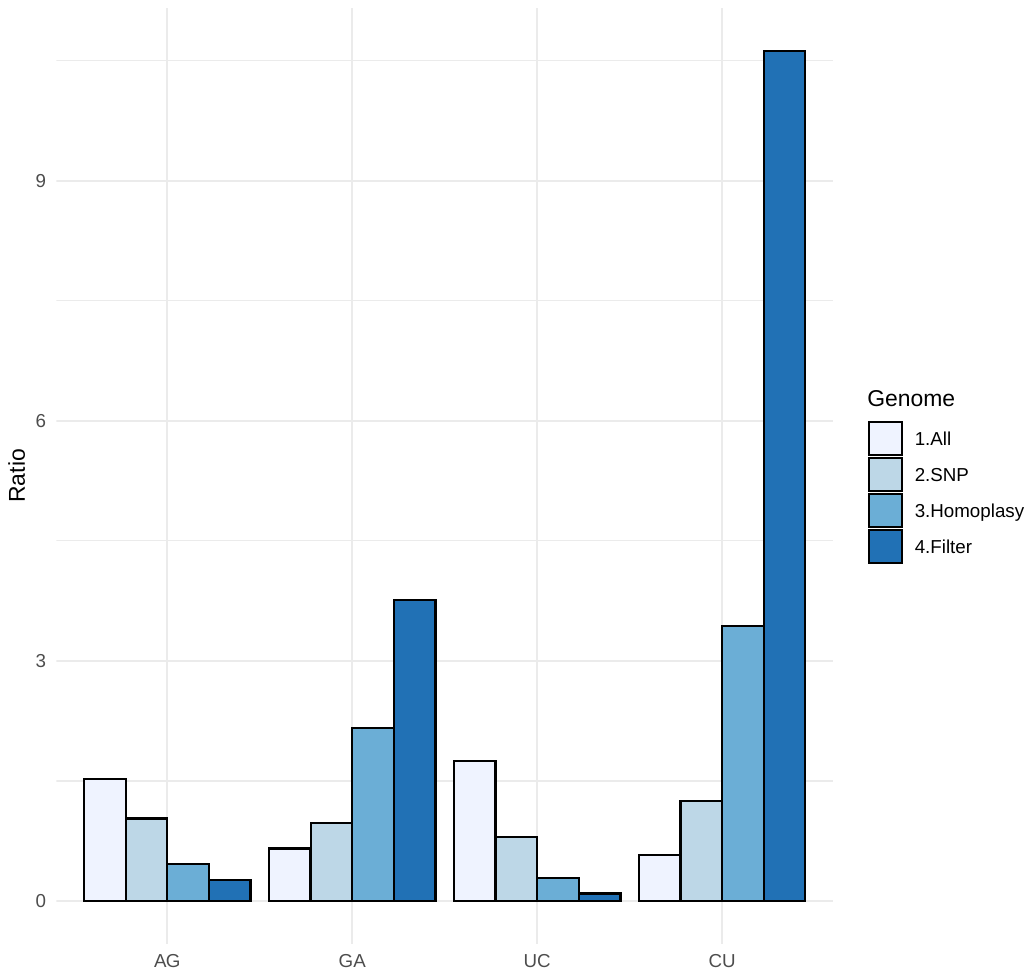

**Figure S6:** Ratio of observed bases (AG:A/G, GA:G/A, UC:T/C, CU:C.T) genome-wide, for solely SNP sites, for homoplasic positions and for filtered homoplasies.

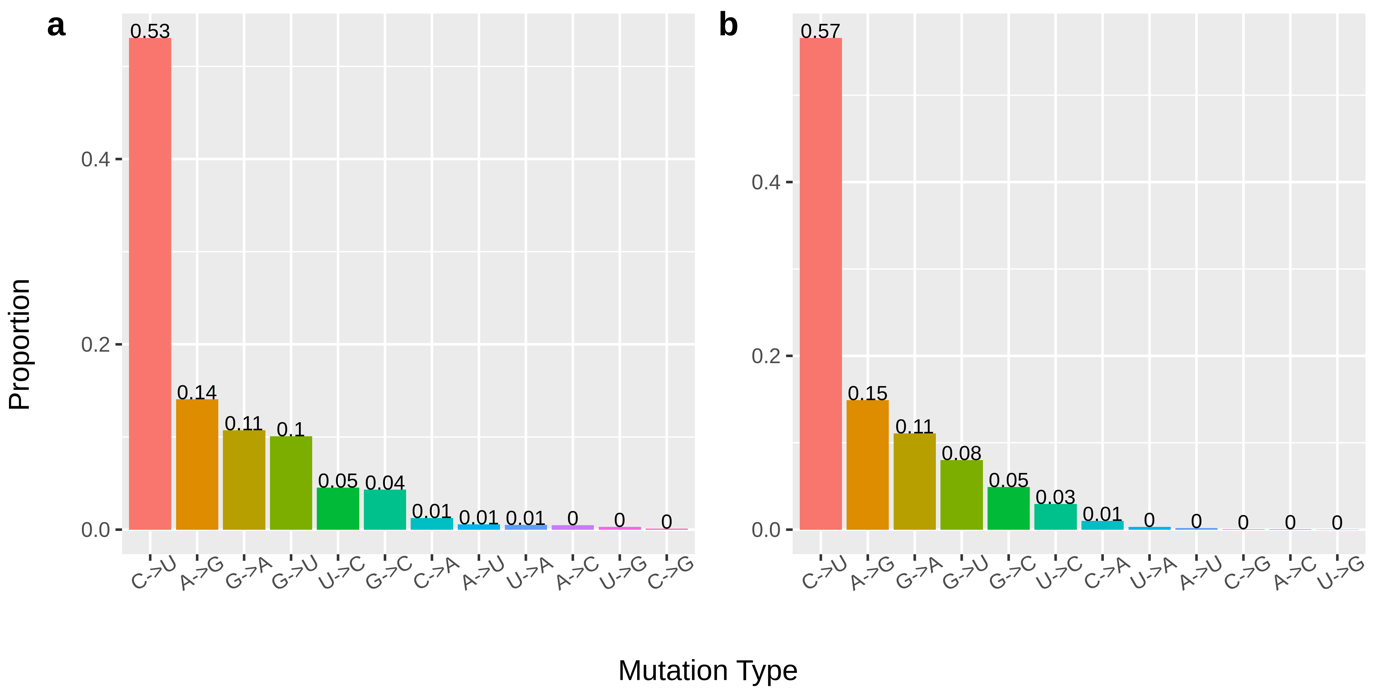

**Figure S7**. Bar plots of the cumulative frequencies of each type of SNP (A) across the entire SARS-CoV-2 genome and (B) across the 308 filtered homoplasic sites. Proportions were computed by dividing frequency of each SNP by the total number of SNPs across all genomes.

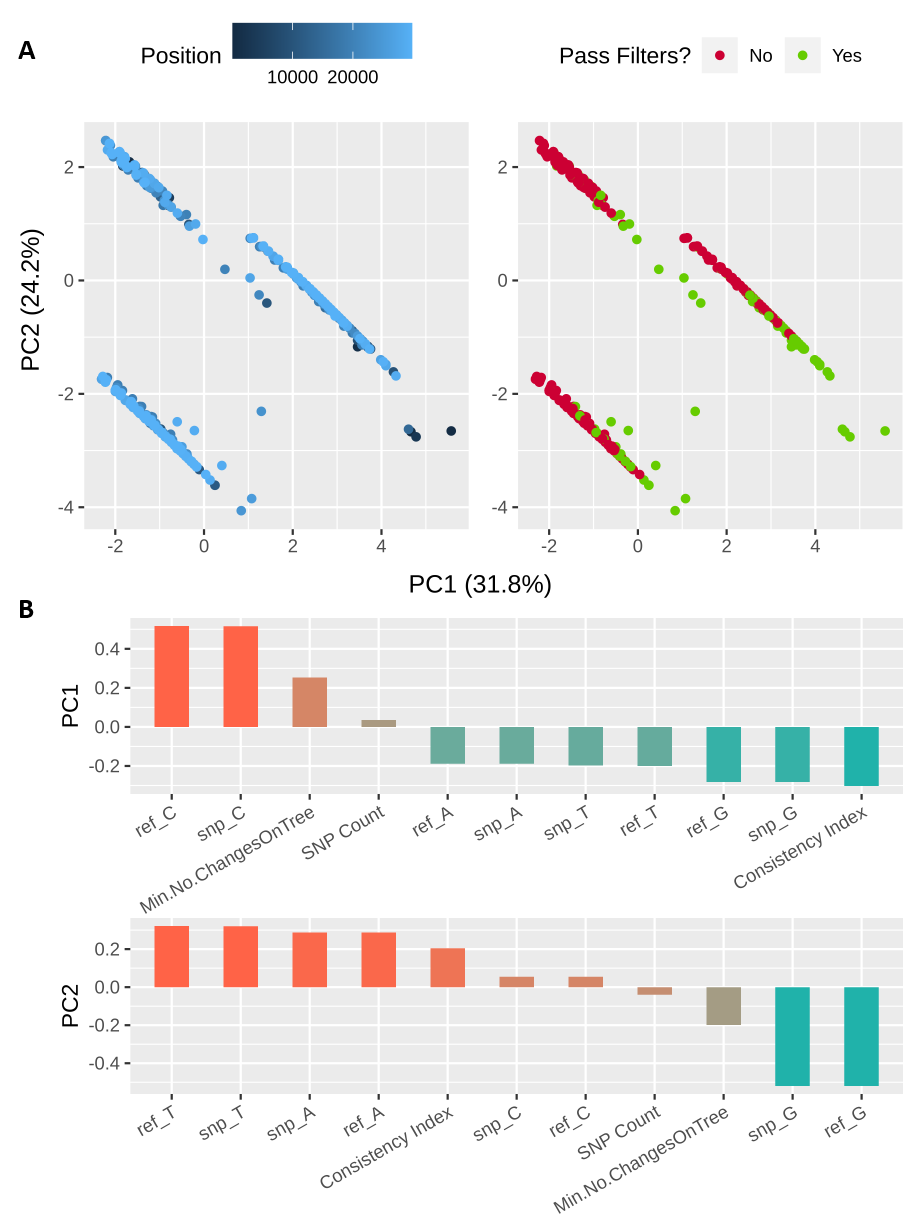

**Figure S8.** (A) Projections of principle components 1 and 2, coloured by genome position on the Wuhan-Hu-1 reference, or by whether the homoplasies were retained after filtering. (B) Loadings of each input variable for principle components 1 and 2.

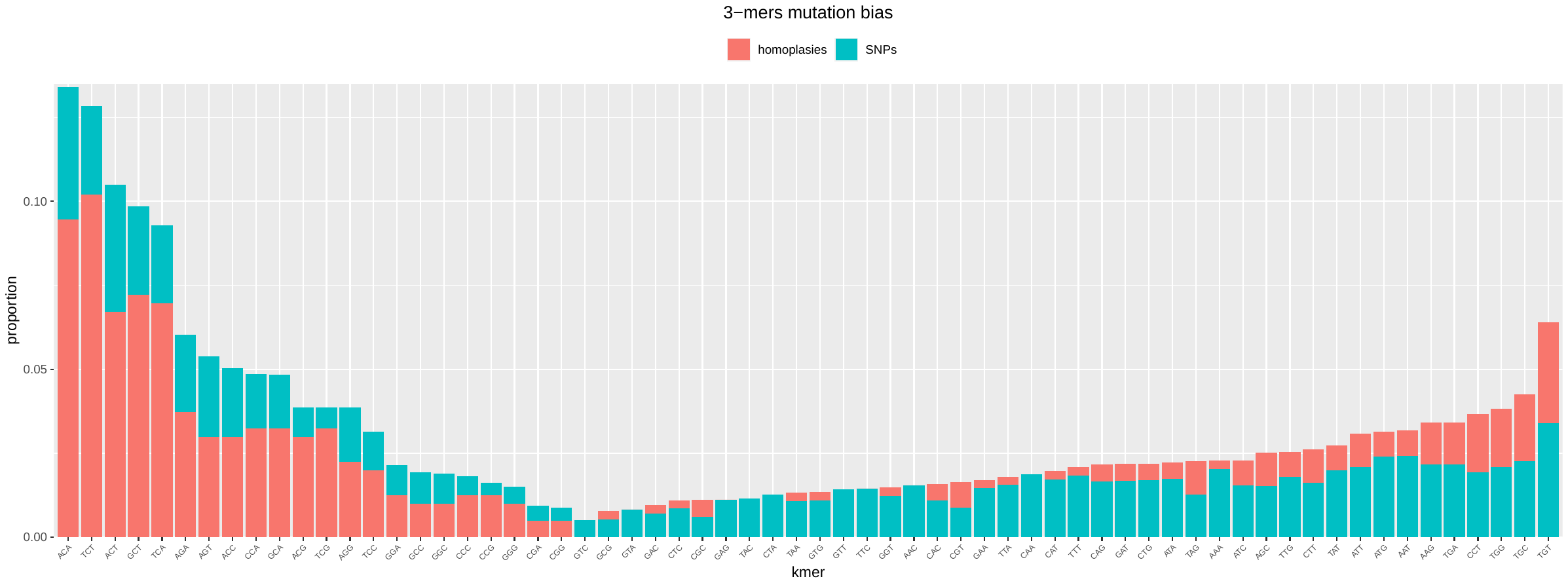

**Figure S9:** Proportion of 3-mers in referent SARS-CoV-2 genome containing a variable base in their central position. The proportions are coloured by filtered homoplastic status and given as the frequency relative to other sites in that state.

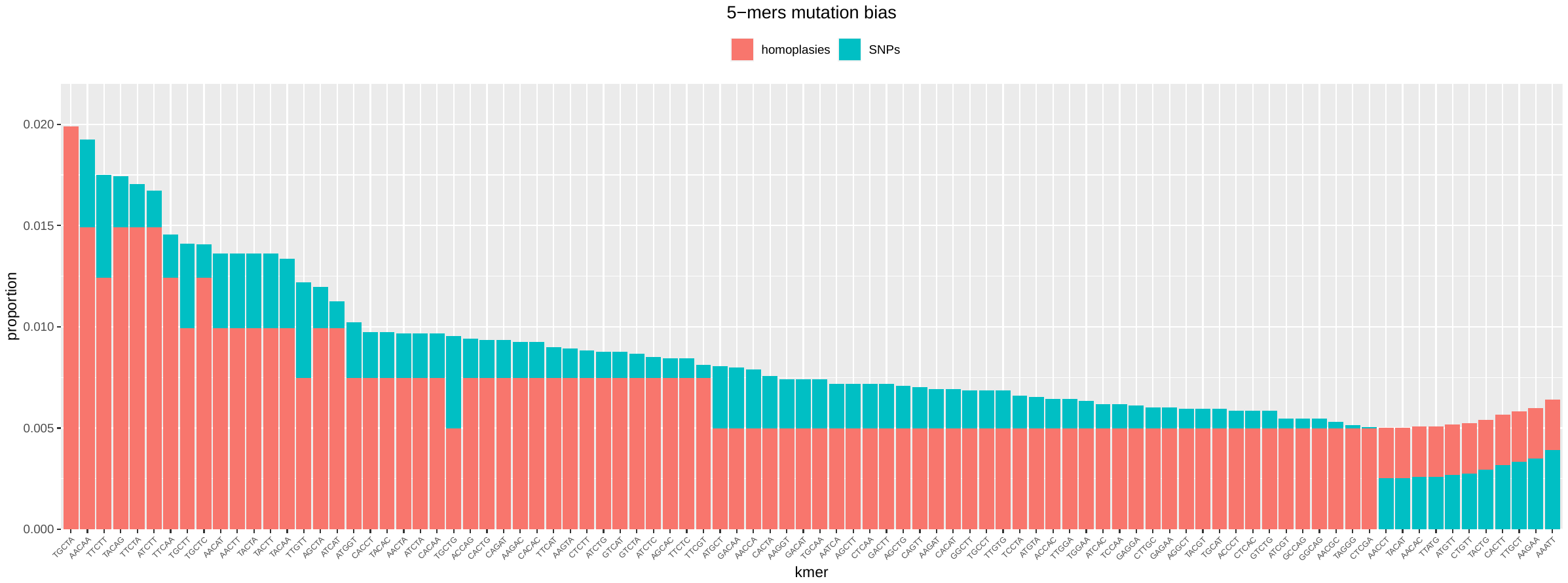

**Figure S10:** Proportion of 5-mers in referent SARS-CoV-2 genome containing a variable base in their central position. The proportions are coloured by the filtered homoplastic status and given as the frequency relative to other sites in that state. Sites with frequency lower than 0.005 were excluded from the figure.

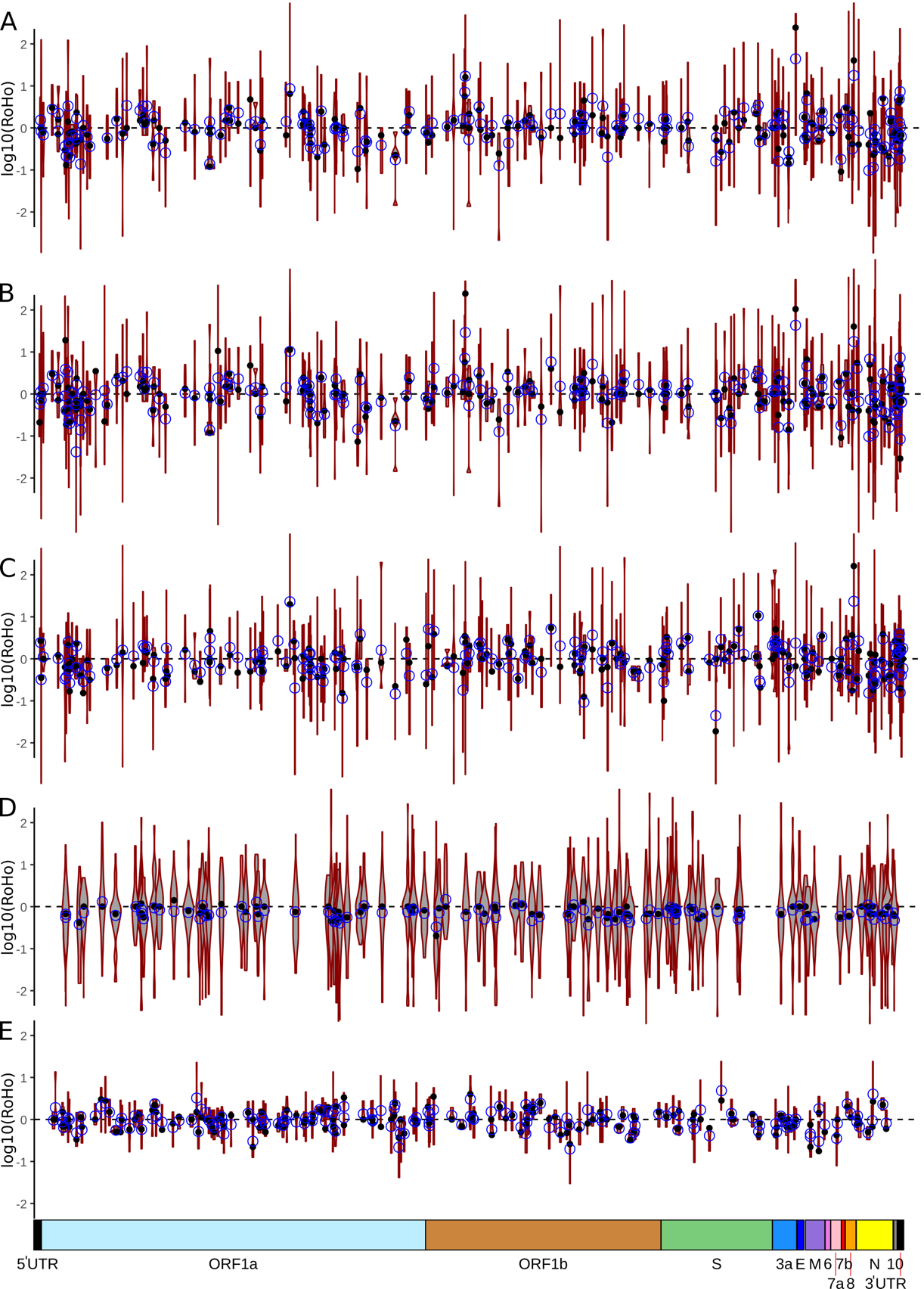

**Figure S11.** Genome-wide Ratio of Homoplasic Offspring (RoHO) scores for homoplasic mutations for which we only enforced a lower minimal number of three independent emergences in the phylogeny. Unless specified otherwise, clades including a secondary homoplasic emergence for the same allele, or having fewer than two descendant tips of each allele are discarded. Masked (de Maio *et al*.) dataset (A); masked (de Maio *et al.*) dataset without discarding embedded homoplasic emergence (B); NextStrain masked dataset (C); 100 randomly generated discrete traits simulated onto the true maximum likelihood phylogeny (D); simulated 10,000 nucleotide alignment of 500 isolates using a 6E-4 substitution rate, see Methods (E). Black dot: median log10(RoHO) value; blue circle: mean log10(RoHO) value. Bottom coloured boxes correspond to encoded ORFs on Wuhan-Hu-1 reference genome.

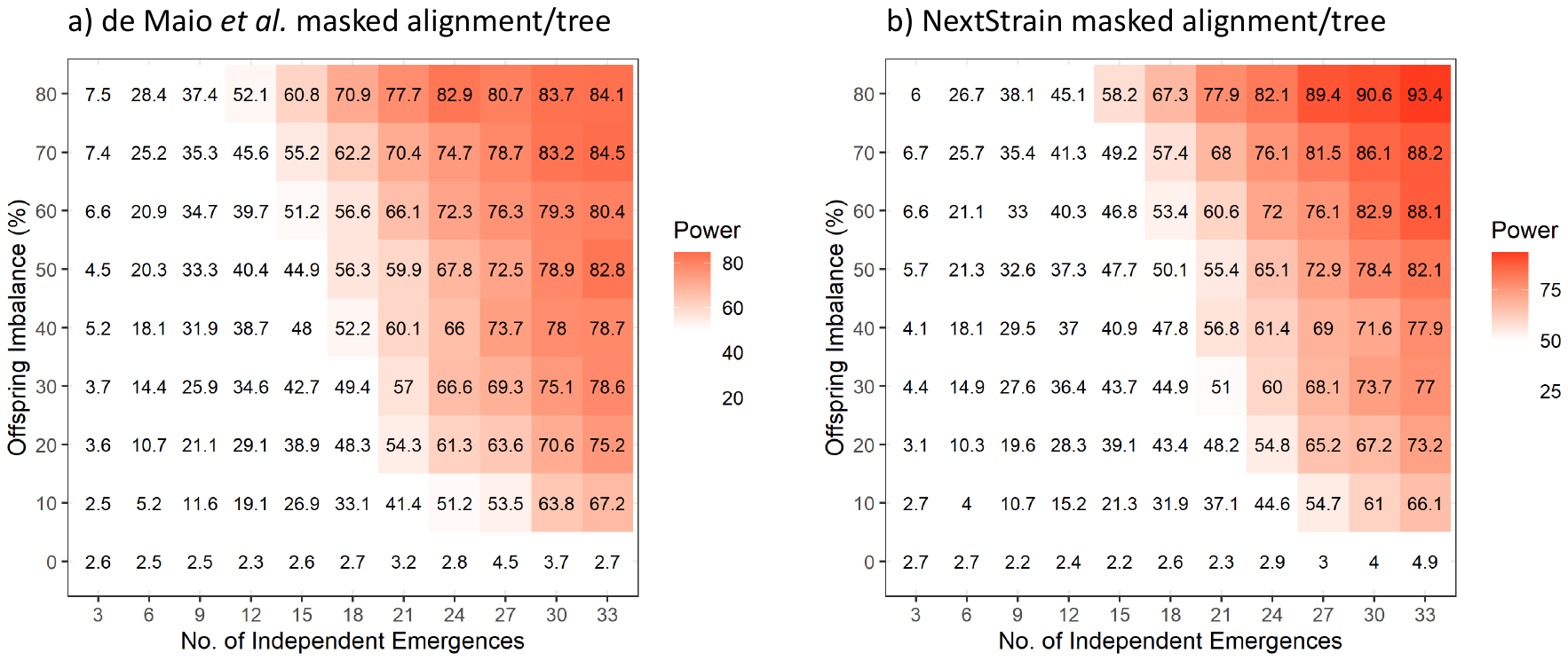

**Figure S12.** Estimation of the detection power of the paired t-tests applied to the RoHO scores

assessed through simulations. Numbers in each box provide the percentage of significant paired t-tests (alpha = 0.05) for 1000 replicates for each combination of 3-33 independent homoplasic emergences and an imbalance in the number of offspring carrying either allele between 0-80% (see Methods). a) Provides results for the de Maio *et al*. masking and b) for the NextStrain masking. A list of masked sites is provided in **Supplementary Table S5**.

**Table S1:** External spreadsheet. List of included GISAID accessions together with full metadata and originating and submitting labs. Those accessions excluded from the original download are provided with justifications.

**Table S2:** External spreadsheet. Variable positions with synonymous and nonsynonymous annotations across the 46,723 assembly alignment used for the homoplasy detection analysis.

**Table S3**: External spreadsheet. Quality metrics of the filtered homoplasies detected in each masked GISAID dataset, see Table S5 for masked positions. Note: positions not annotated as high quality either in the masked or in the alternatively masked GISAID dataset do not appear in the table.

**Table S4**: External spreadsheet. Sheet 1: Metrics associated with the Ratio of Homoplasic Offspring (RoHO) scores for 185 homoplasies. These 185 homoplasies were selected from the 398 homoplasies detected in the masked (de Maio *et al*. 2020) dataset based on phylogenetics parameters (see Methods). Sheet 2: Metrics associated with Ratio of Homoplasic Offspring (RoHO) scores for 199 homoplasies. The 199 homoplasies were selected from the 411 homoplasies detected in the NextStrain masked dataset based on phylogenetics parameters (see Methods).

**Table S5:** Sites masked from the alignment following the masking strategy suggested by de Maio *et al*. <http://virological.org/t/issues-with-sars-cov-2-sequencing-data/473> (focused on masking putative sequencing errors), time stamped to 30/07/2020 and following the NextStrain masking criteria. Unless otherwise stated, masked suggested by de Maio *et al.* was used throughout.

| **Position** | **de Maio *et al*. masking** | **NextStrain masking** |
| --- | --- | --- |
| Beginning of alignment | First 55 | First 130 |
| Singleton sites | 150 153 635 1895 2091 2094 2198 2604 3145 3564 3639 3778 4050 5011 5257 5736 5743 5744 6167 6255 6869 8022 8026 8790 8827 8828 9039 10129 10239 11074 11083 11535 13402 13408 13476 13571 14277 15435 15922 16290 16887 19298 19484 19548 20056 20123 21550 21551 21575 22335 22516 22521 22661 22802 24389 24390 24622 24933 25202 25381 26549 27784 28253 28985 29037 29039 29425 29553 | 18529 29849 29851 29853 |
| End of alignment | Last 100 | Last 50 |

**Table S6:** External excel document. List of homopolymer regions with coordinates and region length identified in the reference genome Wuhan-Hu-1 (GenBank NC_045512.2, GISAID EPI_ISL_402125).
